# Rac1 controls cell turnover and mammary gland reversibility in post-partum involution

**DOI:** 10.1101/2022.02.28.482219

**Authors:** Aleksander Mironov, Matthew Fisher, Randa Elsayed, Melis Karabulutoglu, Nasreen Akhtar

## Abstract

Cell turnover in adult tissues is essential for maintaining tissue homeostasis over a lifespan and for inducing the morphological changes associated with the reproductive cycle. However, the underlying mechanisms that coordinate the balance of cell death and proliferation remain unsolved. Using the mammary gland we have discovered that Rac1 acts as a nexus to control cell turnover. Post-lactational tissue regression is characterized by the death of milk secreting alveoli, but the process is reversible within the first 48h if feeding recommences. In mice lacking epithelial Rac1, alveolar regression was delayed. This defect did not result from failed cell death but rather increased cell turnover. Fitter progenitor cells inappropriately divided, regenerating the alveoli but cell death also concomitantly accelerated. We discovered that progenitor cell hyperproliferation was linked to non-autonomous effects of Rac1 deletion on the macrophageal niche with heightened inflammation. Moreover, loss of Rac1 impaired cell death with autophagy but switched the cell death route to apoptosis. Finally, mammary gland reversibility failed in the absence of Rac1 as the regenerated alveoli failed to recommence lactation upon re-suckling.

## Introduction

Cell turnover in adult tissues is characterized by the death of older cells and replacement with new through stem and progenitor cell proliferation. How these processes are balanced to maintain long-term tissue homeostasis is not clearly understood. The mammary gland is an example of a tissue that maintains a state of flux throughout the adult life. It also undergoes periods of profound growth and regression in each reproductive cycle, providing a tractable model to study cell turnover. The primary role of the mammary gland is to produce milk as a source of nutrients to feed the newborn. Successful lactation depends on the coordinated development and differentiation of the secretory alveolar epithelium during pregnancy and the subsequent removal of these milk-producing units once the milk supply is no longer needed. The balance of cell death and cell division continuously alters to permit tissue growth and regression within the mammary gland reproductive cycle but little is known about how this is coordinated.

Weaning of the infants triggers the mammary gland to enter post-lactational involution, a process in which the surplus milk secreting alveolar epithelium is pruned from the ductal tree using a controlled cell death program (1). In rodents, 90%of the alveolar epithelium laid down in pregnancy is subsequently removed during involution, with the balance of cell death exceeding cell proliferation and the remodeling process completed within approximately two weeks. In dairy animals and humans, the involution process is slower, although cell death occurs, much of the alveolar structure is maintained, either through possible dedifferentiation of existing cells or through cell turnover (2, 3). In murine models of forced involution, simultaneous weaning of the pups at the peak of lactation causes the secretory alveoli to become engorged with milk as production continues for the first 24h, after which they de-differentiate. Both mechanical stretch and milk factors have been reported to stimulate cell death(1, 4). A number of programmed cell death mechanisms have been identified in post-lactational involution, including apoptosis, lysosomal permeabilisation and cell death with autophagy, although the significance of the different death pathways is unclear (5–7).

Moreover, lysosomal permeabilisation and autophagy may feed into an apoptotic death downstream. Involution occurs in two phases; In the first 48h, cell death is triggered but the process of involution is reversible and the gland can re-initiate lactation once suckling resumes (5, 8). The second phase is irreversible and is characterized by destruction of the subtending basement membrane, extensive alveolar cell death and repopulation of stromal adipocytes. The dead cells and residual milk are primarily removed by neighbouring live alveolar mammary epithelial cells (MEC) that act as phagocytes (9–12). Engulfment of milk fat by the non-professional MECs triggers lysosomal permeabilisation through a stat-3 dependent mechanism, which ultimately kills the cells (12). In the second phase, professional phagocytes from the immune system enter the gland to engulf the remaining dead cells and the tissue remodels back to a state closely resembling the nulliparous gland (13, 14).

We previously showed that the Rac1 GTPase plays a crucial role in post-lactational mammary gland remodeling(9). Rac1 is central to the conversion of mammary epithelial cells into non-professional phagocytes for the removal of dead cells and surplus milk and in controlling the inflammatory signature. We now reveal completely new roles for Rac1 in controlling the involution process. We have discovered that Rac1 acts as a nexus, controlling both the rate and balance of cell death and progenitor cell division in involution. Without Rac1 cell turnover accelerates with major consequences on mammary gland reversibility in the first phase of involution. Rac1 therefore has multifaceted roles in orchestrating the involution process.

## Results

### Delayed alveolar regression and repopulation of adipocytes in involuting Rac1-/- mammary glands

Mammary gland remodeling in involution is accompanied by alveolar regression and concomitant fat pad repopulation in the surrounding parenchyma. To investigate the role of Rac1 in tissue regression during involution we examined mammary tissues from female mice triggered to involute through simultaneous weaning of the pups. The Rac1 gene was deleted specifically in luminal cells of the mammary gland using *Rac1^fl/fl^:LSLYFP:WAPiCre* (*Rac1-/-*) conditional knockout mice previously generated (15). Cre negative *Rac1^fl/fl^:LSLYFP* littermates were used as WT controls (WT). Analysis of the WT mammary glands by histology revealed significant lobular alveolar regression and adipocyte repopulation between involution days 2-4 (Fig 1A,C,E,G,I and Fig S1). By contrast alveoli in *Rac1-/-* glands remained distended with virtually no adipocyte repopulation (Fig 1B,D,F,H,I and Fig S1). Four weeks post-involution the alveoli had completely regressed in both WT and *Rac1-/-* glands confirming that lobular alveolar cell death had occurred in the absence of Rac1 (Fig 1J).

**Figure 1:**
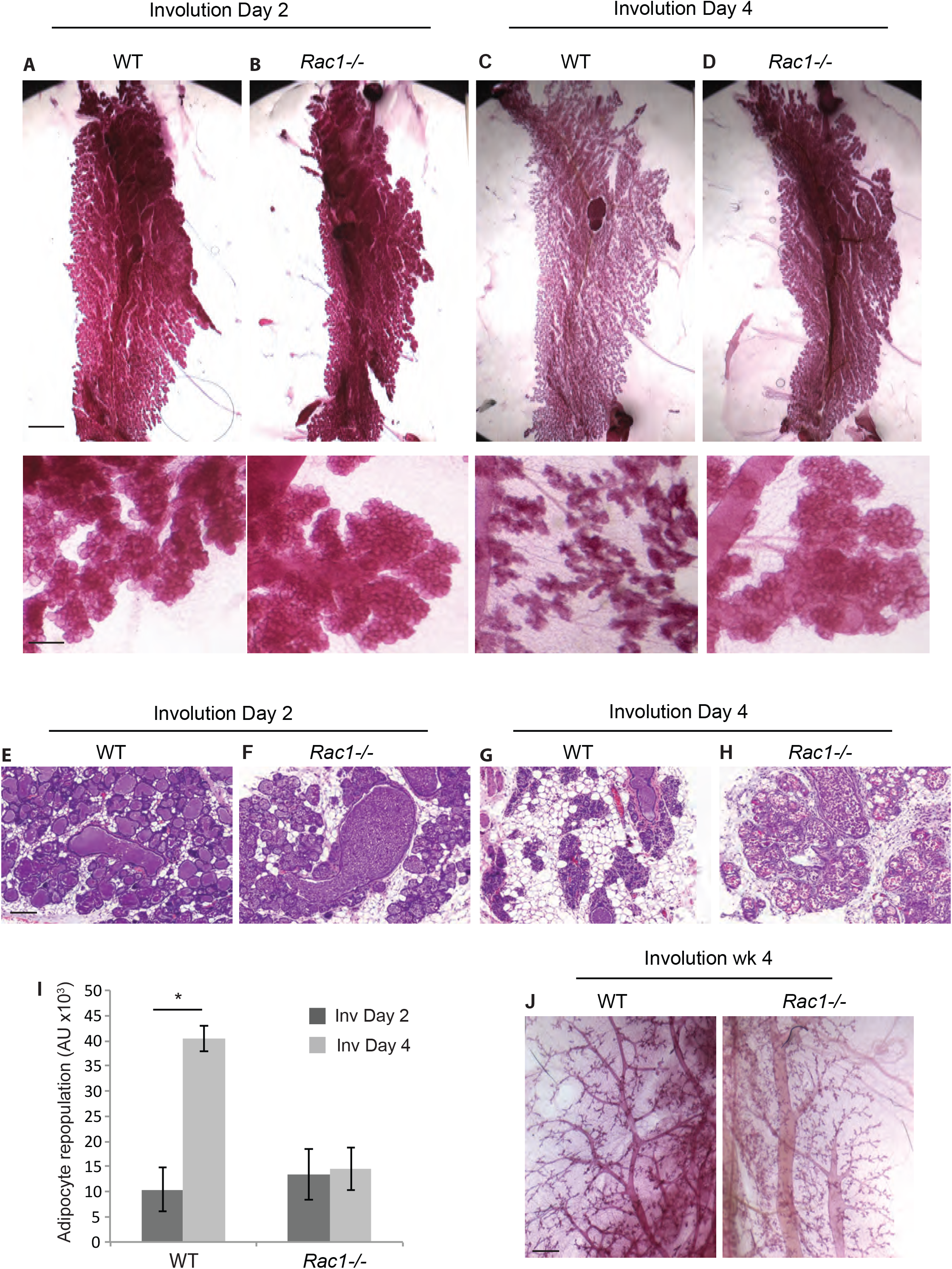
Delayed alveolar regression in Rac1 null glands in involution. (A-D) Carmine staining of whole-mounted mammary glands of WT (A,C) and *Rac1-/-* (B,D) mice at post-lactational involution day 2 and day 4. Note the alveolar regression in WT but not *Rac1-/-* glands at Involution day 4. Bar; 5mm, (inset 1mm). (E-H) Haematoxylin and Eosin (H+E) stain shows alveolar regression and adipocyte repopulation in WT glands (E,G) but this is delayed in *Rac1-/-* glands (F,H). Bar; 100μm. (I) Quantification of adipocyte repopulation. Error bars; +/- SEM of n=3 mice. * P= 0.0043 (J) Carmine staining of whole-mounted WT and *Rac1-/-* mammary glands at 4 weeks post-lactational involution show complete regression of alveoli. Note; bloated ducts persist in *Rac1-/-* glands. Bar; 0.7mm.

### β1-integrin is not upstream of Rac1 in alveolar regression

In mammary gland tissue, β1-integrin functions upstream of Rac1 to regulate lactational differentiation, stem cell renewal and cell cycle progression (9, 16–19), we thus investigated whether β1-integrin was linked to alveolar regression in involution. The β1-integrin gene was deleted in luminal cells of the mammary gland using *β1-integrin^fl/fl^:LSLYFP:WapiCre* conditional knockout mice. Crucially, tissue analysis at involution days 2 and 4 revealed that loss of β1-integrin did not impair alveolar regression and adipocyte repopulation compared to WT mice of a Cre negative genotype (*β1-integrin^fl/fl^:LSLYFP*; Figure 2A-E). This indicates that Rac1’s role in tissue regression is elicited through a distinct upstream signaling axis.

**Figure 2:**
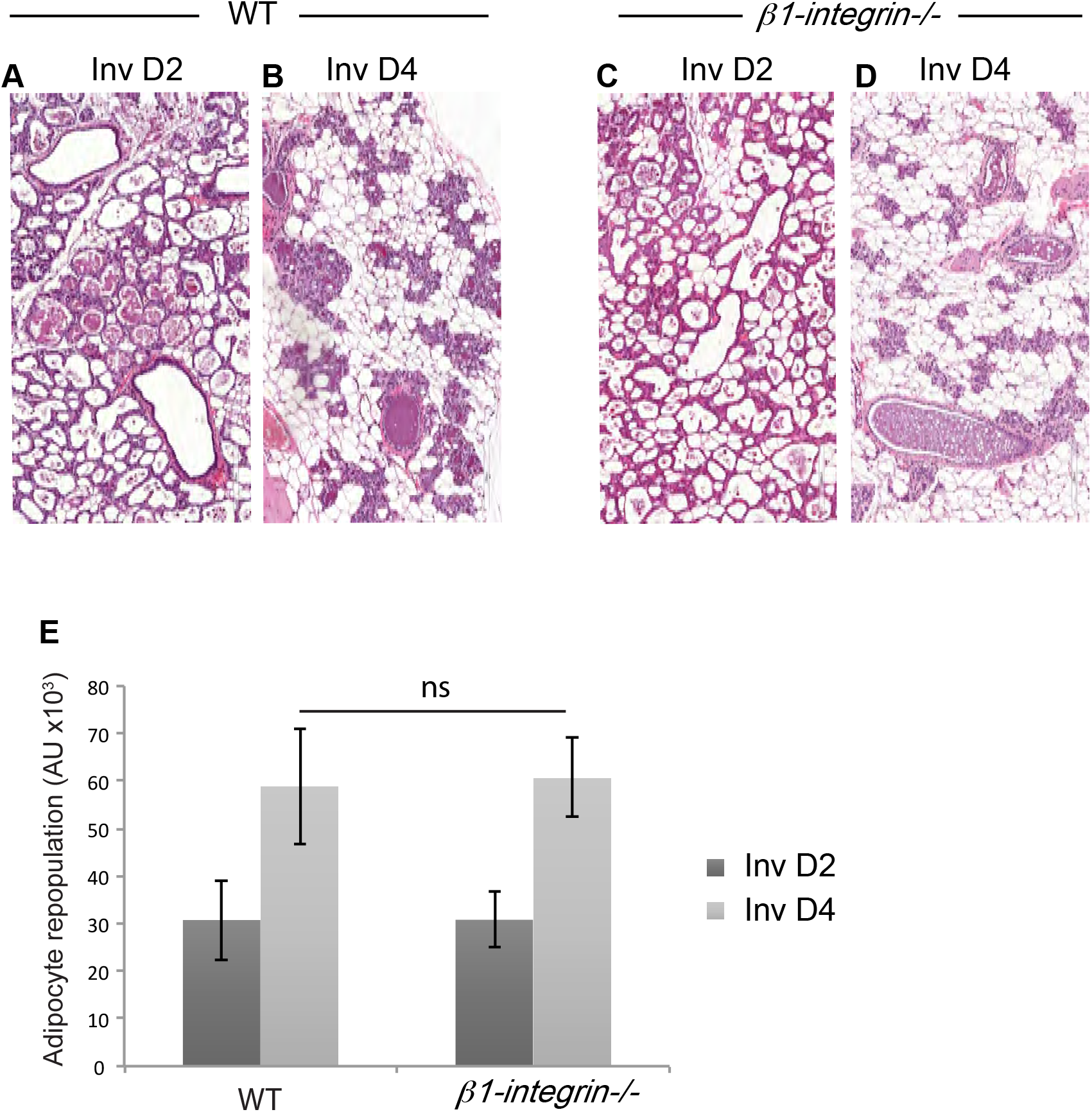
Ablation of β1-integrin does not phenocopy the Rac1-/- involution phenotype. (a-d) H+E stains at Involution day 2 and 4 show no delay in alveolar regression in *β1-integrin-/-* mice. Both WT glands (a,b) and *β1-integrin-/-* (c,d) glands equally regressed. Bar: 100μm. (e) Quantification of adipocyte repopulation. Error bars; +/- SEM of n=4 mice

### Rac1 ablation imbalances cell turnover rates in involution causing delayed alveolar regression

For effective removal of surplus alveoli the balance of cell death should exceed cell proliferation. We first investigated whether there was an initial delay in cell death in *Rac1-/-* glands, which might explain the delay in alveolar regression. At involution day 4 where the delayed regression is most prominent, numerous cell corpses were evident in *Rac1-/-* glands both within the alveolar epithelium and within the lumen space (Figure 3A). We confirmed the dead cells with cleaved caspase −3 staining and that they were of luminal cell origin with the *Rosa:LSL-YFP* reporter gene which is activated in response to WAPiCre induced recombination (Figure 3B,C). Moreover, cell death was not delayed at earlier involution stages (day 2), but rather increased cell corpses were detected in *Rac1-/-* tissue lumens (Figure 3D,E). However, this might be a result of defective clearance by epithelial phagocytes as opposed to increased cell death in *Rac1-/-* glands (9). Thus the delay in alveolar regression in transgenic glands is not linked to impaired cell death.

**Figure 3:**
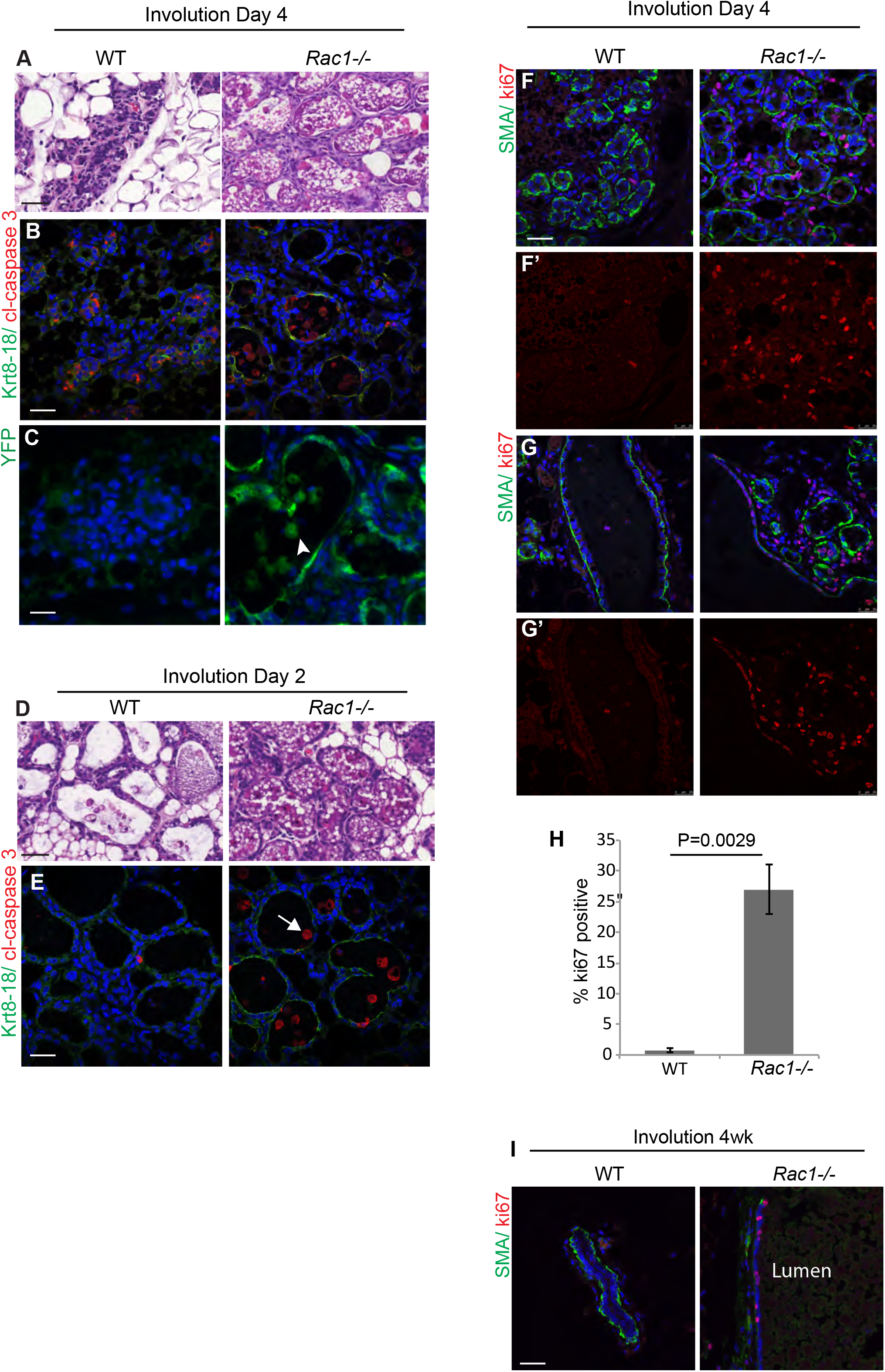
Delayed alveolar regression is not due to impaired cell death but heightened proliferation. (A,D) H+E stain shows numerous dead cells in *Rac1-/-* alveolar lumens at Involution day 4 (A) and Involution day 2 (D). Bar; 40μM. (B,E) Dead cells were confirmed in involution day 4 (B) and day 2 (E) glands by immunofluorescence staining with cleaved caspase 3 (red) and Keratin (Krt) 8/18 (green) to mark luminal cells. Note the Krt8-18 antibody cross reacts with intact keratins in live cells and not cleaved forms in dead cells. Arrow: caspase positive dead cells in the lumen. Bar; 40μm. (C) Dead cells in the lumen were confirmed to be of luminal origin with the YFP reporter gene. GFP antibody was used to stain the YFP reporter gene. Arrowhead: dead cells in the lumen. Bar; 20μm. (F,G) Immunofluorescence staining for proliferation marker Ki67 (red) reveals increased proliferation within *Rac1-/-* glands at involution day 4 in alveoli (F, F’) and ducts (G, G’). Epithelial tissue boundary was detected by smooth muscle actin (SMA, green) present in myoepithelial cells. Bar; 50μm. (H) Quantitative analysis of Ki67 staining at involution day 4. 10 areas/mouse were analysed. Error bars: +/- SEM of n=3 mice. P=0.0029. (I) Proliferation persists in bloated *Rac1-/-* ducts 4 weeks postlactational-involution. Bar; 50μm.

Retention of milk within the lumens might contribute to the distended alveolar phenotype in *Rac1-/-* glands, since Rac1 is crucial for epithelial cell-directed engulfment of apoptotic cell corpses and residual milk(9). However, given the extensive cell death detected in early involution it was surprising that the alveoli remained intact. We thus investigated whether *Rac1-/-* alveoli were being maintained through cell renewal. Ki67 staining revealed almost no detectable proliferation in WT glands at involution day 4. In contrast *Rac1-/-* mammary glands showed extensive proliferation in both the ducts and alveoli (Figure 3F-H and Figure S2). Proliferation within the transgenic ducts continued at 4 weeks post-involution, at this stage however, most of the lobular alveoli had regressed (Figure 3I and 1J). Taken together these results suggest that the delayed alveolar regression in early involution in *Rac1-/-* glands is linked to increased compensatory cell proliferation and not delayed apoptosis. Thus Rac1 has a key role in balancing the rate of cell death and proliferation in involution by limiting the division progeny of progenitors. Without Rac1 alveolar progenitors divide unexpectedly in involution. The newly replaced cells however, have a limited lifespan and succumb to death as evidenced by subsequent alveolar regression 4 weeks post-weaning involution.

### Loss of Rac1 elicits distinct proliferation responses within the first and second gestation

Rac1 has been linked to cell cycle progression in numerous cultured cells and *in vivo* tissues(19–22). This is in complete contrast to our findings *in vivo* in the involuting mammary gland where loss of Rac1 induces proliferation. We thus examined the effects of Rac1 deletion on glandular development and proliferation within the first and second gestation. In the first gestation, wholemount analysis of mammary glands at pregnancy day 18 revealed slightly smaller alveoli in *Rac1-/-* glands (Figure 4A). However, immunostaining with Ki67 at lactation day 2 in the first cycle revealed no significant difference in proliferation between WT and *Rac1-/-* glands (Figure 4B,C). Moreover detection of the *Rosa:LSL:YFP* reporter gene revealed that recombination and thereby gene deletion was extensive and the proliferation was occurring within YFP positive/*Rac1-/-* cells and not from cells that had escaped recombination (Fig 4B).

**Figure 4:**
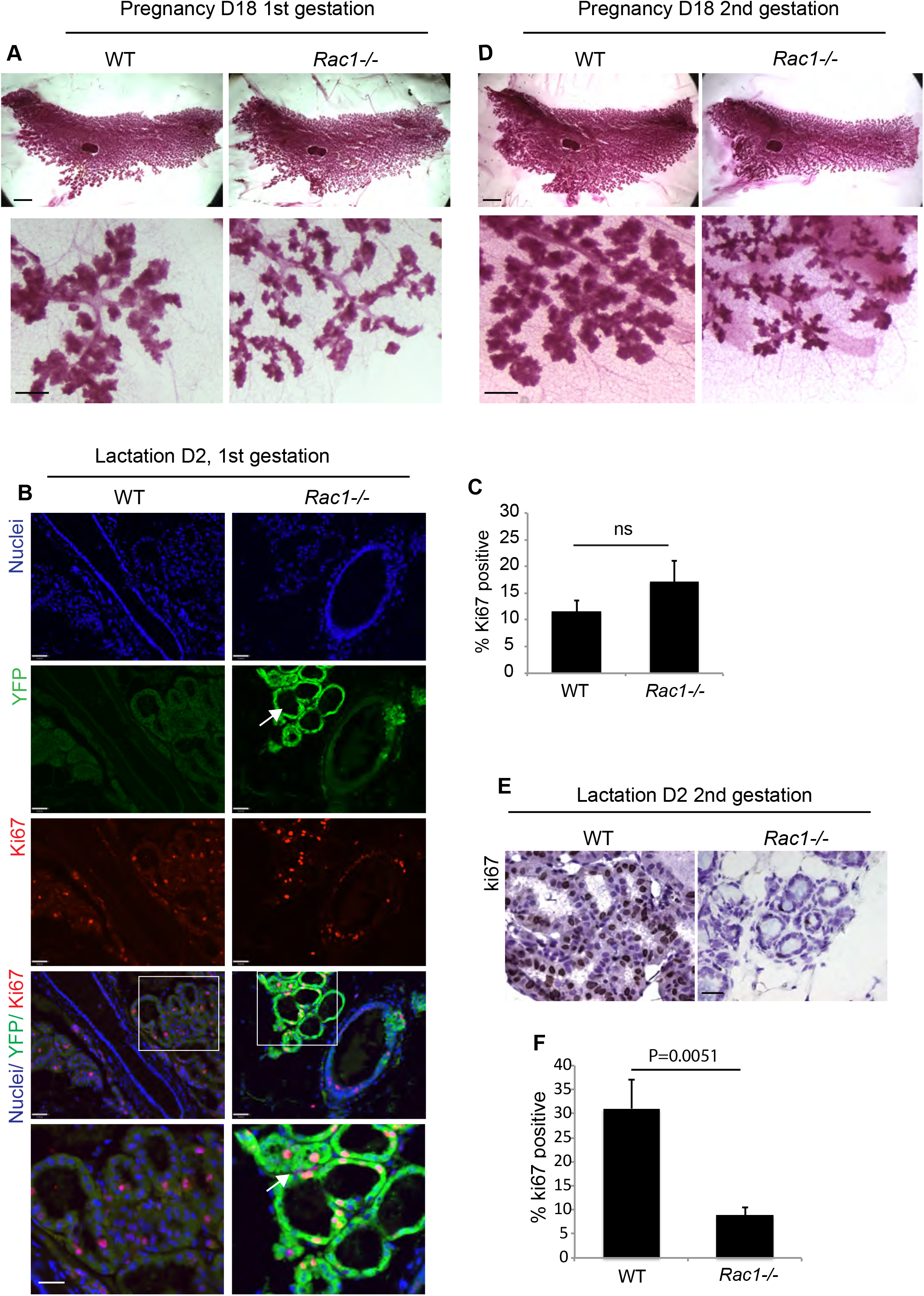
Distinct proliferation in Rac1-/- mammary glands within the first and second gestations. (A) Carmine staining of whole-mounted mammary glands from WT and *Rac1-/-* mice at pregnancy day 18 in the first gestation. Bar; 2.8mm (insert 0.3mm). (B) Immunoflourescence staining with Ki67 in WT and *Rac1-/-* glands shows no difference in proliferation at lactation day 2 following the first gestation. Green fluorescent protein antibody was used to detect the WapiCre driven YFP reporter gene expression and hence Rac1 deletion. Arrows show proliferation in YFP positive/ *Rac1-/-*cells. Bar; 40μm. (C) Quantitative analysis of Ki67 staining at lactation day 2 following the first gestation shows no significant difference between WT and *Rac1-/-* glands. 10 areas/mouse were analysed. Error bars: +/- SEM of n=4 WT mice and n=5 *Rac1-/-* mice. P>0.05. (D) Carmine staining of whole-mounted mammary glands from WT and *Rac1-/-* mice at pregnancy day 18 in the second gestation. Note the reduced lobular alveolar development. Bar; 2.8mm (insert 0.3mm). (E) Ki67 staining reveals reduced proliferation in *Rac1-/-* mammary glands at lactation day 2 following the second gestation. Bar; 40μm. (F) Quantitative analysis of ki67 positive staining in WT and *Rac1-/-* glands at lactation day 2, second gestation. 15 areas/ mouse were analysed. Error bars: +/- SEM of n=4 mice. * P= 0.0051

In marked contrast to the involuting gland, within the second lactation cycle at day 2, there was a severe block in proliferation of luminal cells with concomitant reduced lobular alveolar development in both late pregnancy and early lactation stages (Figure 4D-F, Fig S3). Taken together, these data show that Rac1 deletion has no effect on proliferation within the first lactation, heightened proliferation in post-lactational involution and severely defective proliferation within the second lactation. The disparity in proliferation profiles suggests stage-specific cell autonomous and possible non-autonomous regulation by Rac1.

### Increased proliferation in involuting Rac1-/- glands is linked to inflammatory signals

To explain the contrasting effect on cell proliferation within different stages of the mammary gland cycle, we sought to investigate possible cell non-autonomous effects of Rac1. We previously showed heightened inflammatory responses in *Rac1-/-* mammary glands in post-lactational involution(9). Heightened inflammation has been linked to altered epithelial proliferation and pathogenesis within numerous tissues including the gut and skin we thus sought to investigate whether altered inflammatory responses without Rac1 were linked to the increased proliferation. To test this we first examined for presence of inflammatory signals within in the gestational stage and in involution. In the first gestation, Rac1 deletion did not induce inflammatory signals (Figure 5A, C,D) and this correlates with no significant difference in proliferation (Fig 4B,C). In contrast in involution Rac1 deletion heightens inflammatory signals with early macrophage recruitment (9) (Figure 5B-D) and this correlates with elevated proliferation (Figure 3F-H). To remove possible impending signals from inflammatory cells, pure populations of mammary epithelia were isolated from *Rac1^fl/fl^CreER^TM^* mice and the Rac1 gene was ablated in culture with 4-hydroxytamoxifen. At this stage greater than 90% of the cells are alveolar in origin. Loss of Rac1 impaired proliferation in 3D organoids on a basement membrane matrix (Figure 5E-G). Moreover ductal cultures from nulliparous *Rac1^fl/fl^CreER^TM^* mammary glands failed to branch out in the absence of Rac1 (Figure 5F). Together, these data suggest that increased proliferation in involution in *Rac1-/-* glands is in part driven through secondary non-cell autonomous effects of Rac1 deletion and is connected to heightened and sustained inflammatory responses. These findings reveal that Rac1 mediates a functionally important cross-talk between mammary gland cells and immune phagocytes within their microenvironmental niche.

**Figure 5:**
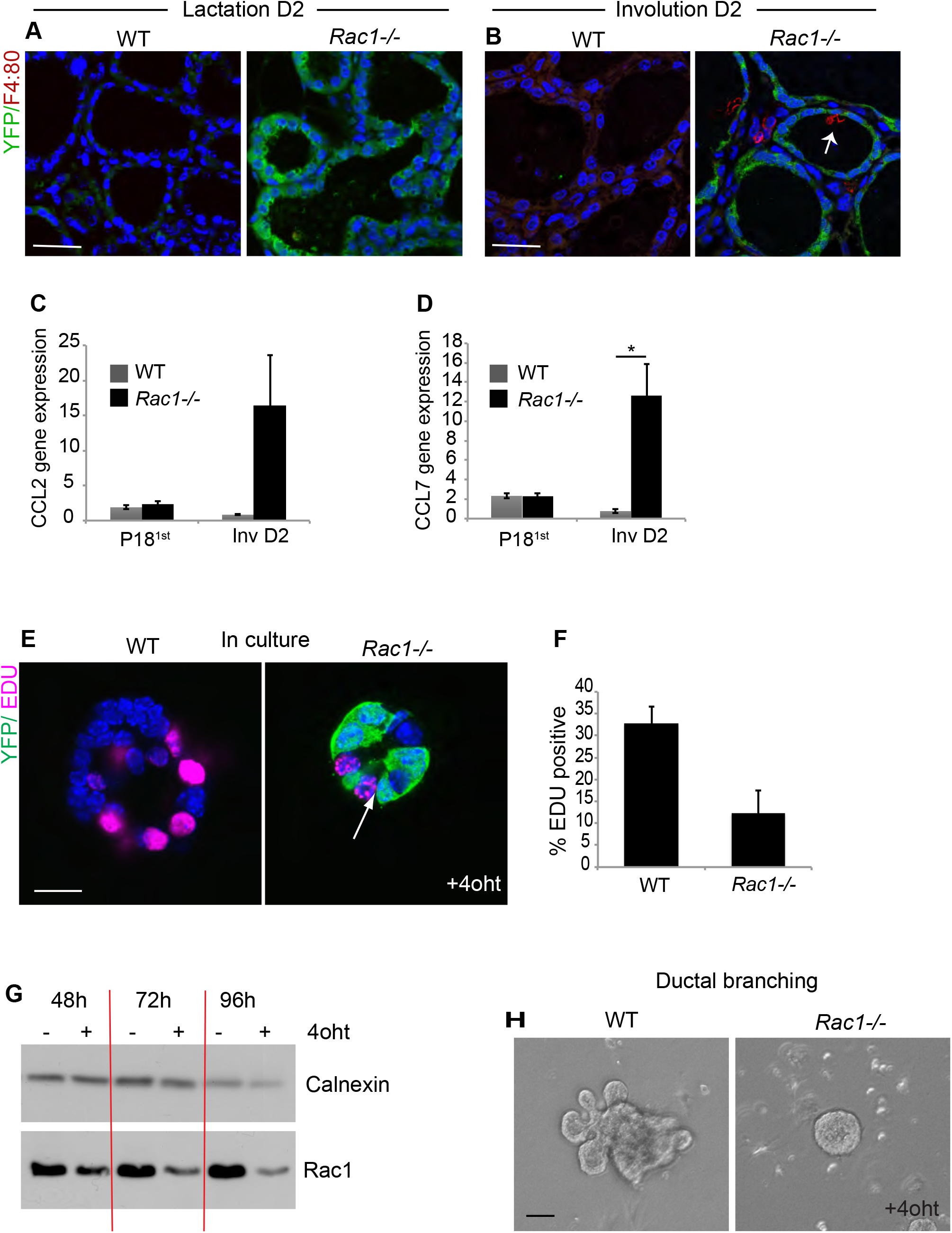
Rac1-/- hyperproliferation in involution is linked to inflammation. (A,B) Immunoflourescence staining with F4:80 antibody (red) to detect macrophages in WT and *Rac1-/-* glands at Lactation day 2 and Involution day 2. GFP antibody was used to detect the YFP reporter gene expression. Error bars: +/- SEM of n=3-4 mice. Arrow; macrophage in lumen of *Rac1-/-* alveoli. Bar; 40μm. (C,D) Quantitative RT-PCR shows elevated inflammatory chemokines CCL2 and CCL7 in *Rac1-/-* glands at Involution day 2 but not in late pregnancy. * P= <0.025. (E) EDU incorporation in WT and *Rac1-/-* primary mammary alveolar organoids from *Rac^flfl^CreERT* mid-pregnant mice, cultured on a BM-matrix show reduced proliferation in the absence of Rac1. Immunofluorescence staining with GFP antibody was used to detect the YFP reporter gene expression (green) and EDU (magenta). Images are confocal sections through the middle of the alveoli. Arrow: proliferation in YFP negative cells in *Rac1-/-* organoid. Bar; 20μm. (F) Quantitative analysis of EDU incorporation reveals decreased proliferation in *Rac1-/-* organoids. EDU incorporation was counted in YFP positive cells/*Rac1-/-* only in 4oht treated organoids. Error bars: +/- SEM of n=3 experiments. (G) Depletion of Rac1 in primary MEC organoids was confirmed by immunoblotting cell lysates prepared from control or 4oht treated cultures with a Rac1 antibody. Calnexin antibody was used to show equal loading of protein. (H) Ductal branching in WT and *Rac1-/-* organoids in response to FGF stimulation. Depletion of Rac1 prevents outgrowth. Bar; 40μm.

### Rac1 directs cell death with autophagy but not apoptosis or necrosis

As cell death and inflammation are intermittently linked, we examined the effects of removing Rac1 on the cell death route in involution. Ultrastructural studies in day 2 and 4, involuting glands revealed numerous vacuolar structures in WT alveolar epithelium but not *Rac1-/-* transgenics (Figure 6A,G,M,O). Further analysis revealed autophagosomes, autolysosomes and lysosomes in the WT alveolar epithelium, suggestive of cell death with autophagy (Figure 6A-C, M,N). Moreover, we detected numerous phagosomes with engulfed milk proteins, milk lipid droplets and dead cells (Figure 6D-F), which supports are previous findings showing engulfment through phagocytic cups and macropinosomes in WT cells(9). In contrast *Rac1-/-* alveoli were completely void of both autophagososmes and phagosomes, instead we detected either live cells or dead cells shed into the lumen with apoptotic morphology and some necrotic cells with ruptured membranes and organelles released extracellularly (Figure 6G-L,O,P). We further confirmed autophagy in WT glands by immunostaining for the essential autophagy related LC3β protein, which appeared punctate and therefore indicative of translocation to the autopagososme membrane. In contrast, LC3β staining was diffuse in *Rac1-/-* transgenics confirming the absence of autophagosomes (Figure 6Q, R). Consistent with the ultrastructural studies, gene array analysis revealed down-regulation of genes associated with autophagososme and lysosomal pathways (Figure 6S). Whole groups of lysosomal hydrolases, including proteases, glycosidases, sulfatases, DNases and lipases were down-regulated in *Rac1-/-* transgenics (Table S1). Taken together these data show that Rac1 is required to induce cell death with autophagy but without Rac1 cells can still die via apoptosis and necrosis. The presence of necrotic cells likely contribute to the heightened inflammatory responses in *Rac1-/-* glands.

**Figure 6:**
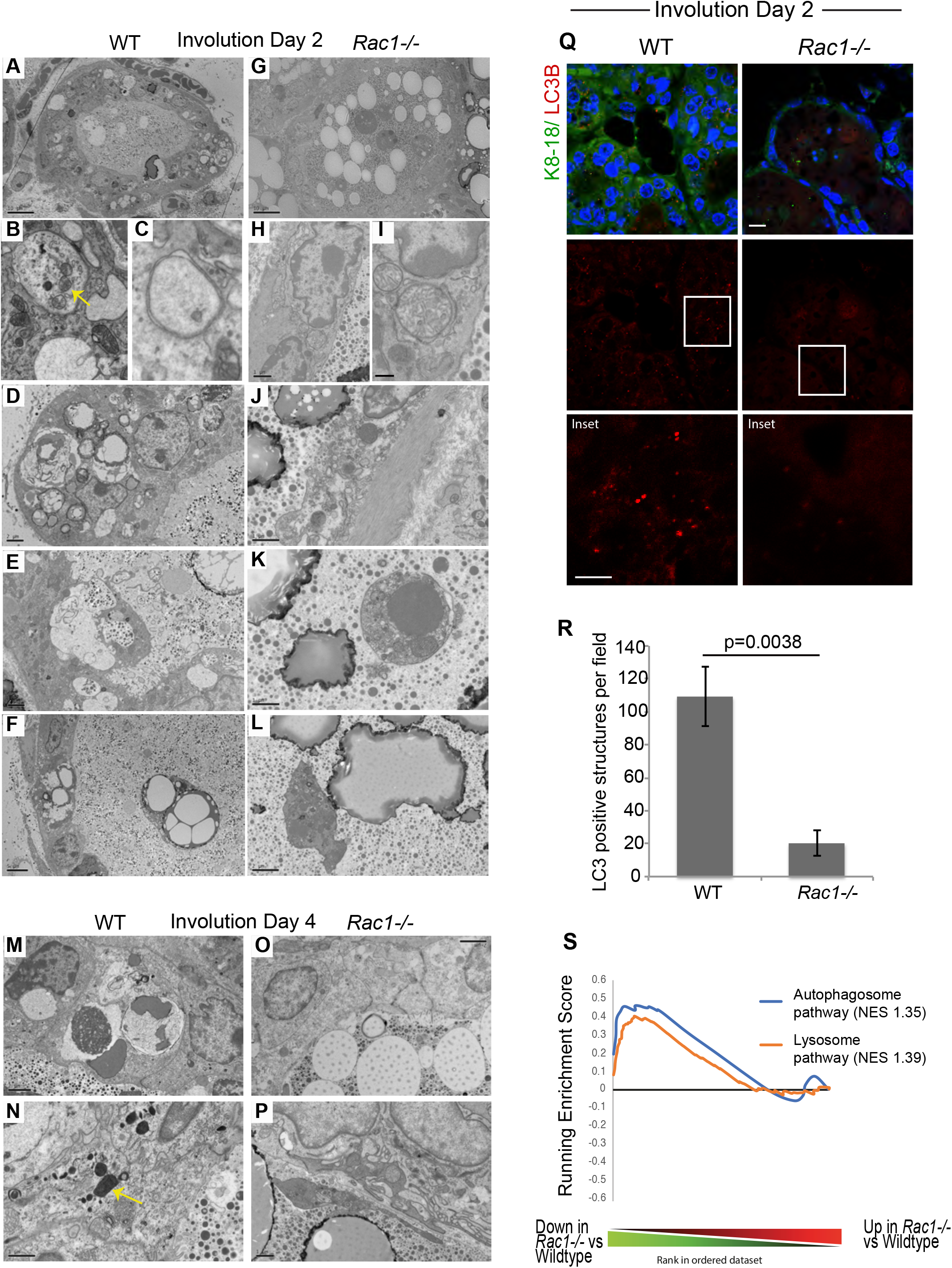
Rac1 mediates cell death with autophagy but not apoptosis. Electron micrographs of WT (A-F) and *Rac1-/-* (G-L) involution day 2 glands. (A,) WT alveolus showing engulfment activity with numerous phagosome-like structures within the epithelium. Bar; 10μM (B,C) Autophagosomes in WT luminal cells; Note mitochondria in autophagosomes (B; arrow). (D) Autophagosmes and phagosomes containing milk lipids in WT cells. Bar; 2μm. (E) Phagosome forming around engulfed milk in WT cells. Bar; 2μm. (F) Lipid droplets associated with WT dead cells within the epithelium and shed into the lumen. Bar; 5μm. (G) *Rac1-/-* alveolus showing milk lipid droplets and dead cells in the lumen but no phagosome like structures within the epithelium. Bar; 10μm. (H,I) No autophagosomes in *Rac1-/-* epithelium. Bar; (H) 1μm, (I) 0.4μm. (J) Cell necrosis in *Rac1-/-* epithelium. Bar; 1μm (K,L) Apoptotic *Rac1-/-* cells shed into the lumen with nuclear pyknosis (K) and membrane blebbing (L). Bar; (K) 1μm, (L) 2μm. (M-P) Involution day 4 WT cells showing autophagososmes, phagosomes (M) and lysosomes (arrow; N) but Rac1-/- (O,P) show none. Bar; (M,O) 2μm, (N,P) 1μm. (Q) LC3β antibody was used to detect autophagic structures in involution day 2 tissues. Note; vesicles in WT epithelium but not *Rac1-/-*. Krt8/18 antibody was used to mark luminal cells. Bar; 10μM (insert 6μm). (R) Quantitative analysis shows markedly reduced LC3β vesicles in *Rac1-/-* cells. LC3 positive structures were counted per field. Error bars: +/- SEM of n=3 mice. P= 0.0038. (S) GSEA demonstrating down-regulation of autophagosome and lysosome genes in *Rac1-/-* glands compared with WT at involution day 2. NES; normalized enrichment score.

### Cell death is accelerated without Rac1

We next investigated whether necrotic cells occurred secondary to apoptosis as a result of defective phagocytosis or whether loss of Rac1 triggers a programmed necrotic cell death. To test this directly, primary cultures of WT and *Rac1-/-* cells were induced to undergo anoikis, a detachment induced cell death. We chose this method of cell death for two reasons, first, to prevent dying cells from removal by neighbouring non-professional phagocytosis, as single cells suspended in media are spatially out of reach for phagocytic removal and second to trigger an innate programmed cell death rather than chemical-induced. Dying cells incorporated higher levels of propidium iodide (PI) in the absence of Rac1 compared to WT controls, suggesting cell death by necrosis or late stage apoptosis (Figure 7A,B). To establish the proximal cell death route, we first examined for hallmarks of apoptosis as this process accompanies a series of well-defined biological steps. Both WT and *Rac1-/-* cell corpses displayed intact membranes with nuclear condensation, late stage membrane blebbing, body fragmentation and stained positive for cleaved caspase 3 indicative of an apoptotic cell death (Figure 7C-G and 3B,E). Early stage apoptosis is characterized by phosphatidylserine exposure to the outer membrane leaflet and Annexin V is commonly used to detect this motif. Co-labelling with annexin V and PI in cells suspended for 1h and 5h revealed that approximately the same number of cells enter apoptosis with and without Rac1, (Annexin V only) however, by 5h significantly more *Rac1-/-* cells proceed to late stage apoptosis/necrosis (Annexin V/ PI) with a concomitant reduction in numbers in early stage apoptosis (Annexin V only; Figure 7H). Moreover, the numbers of viable cells declined following 8h suspension in the absence of Rac1 indicating that cells had proceeded through death and disintegrated within this time frame compared to WT controls (Figure 7I, J).

**Figure 7:**
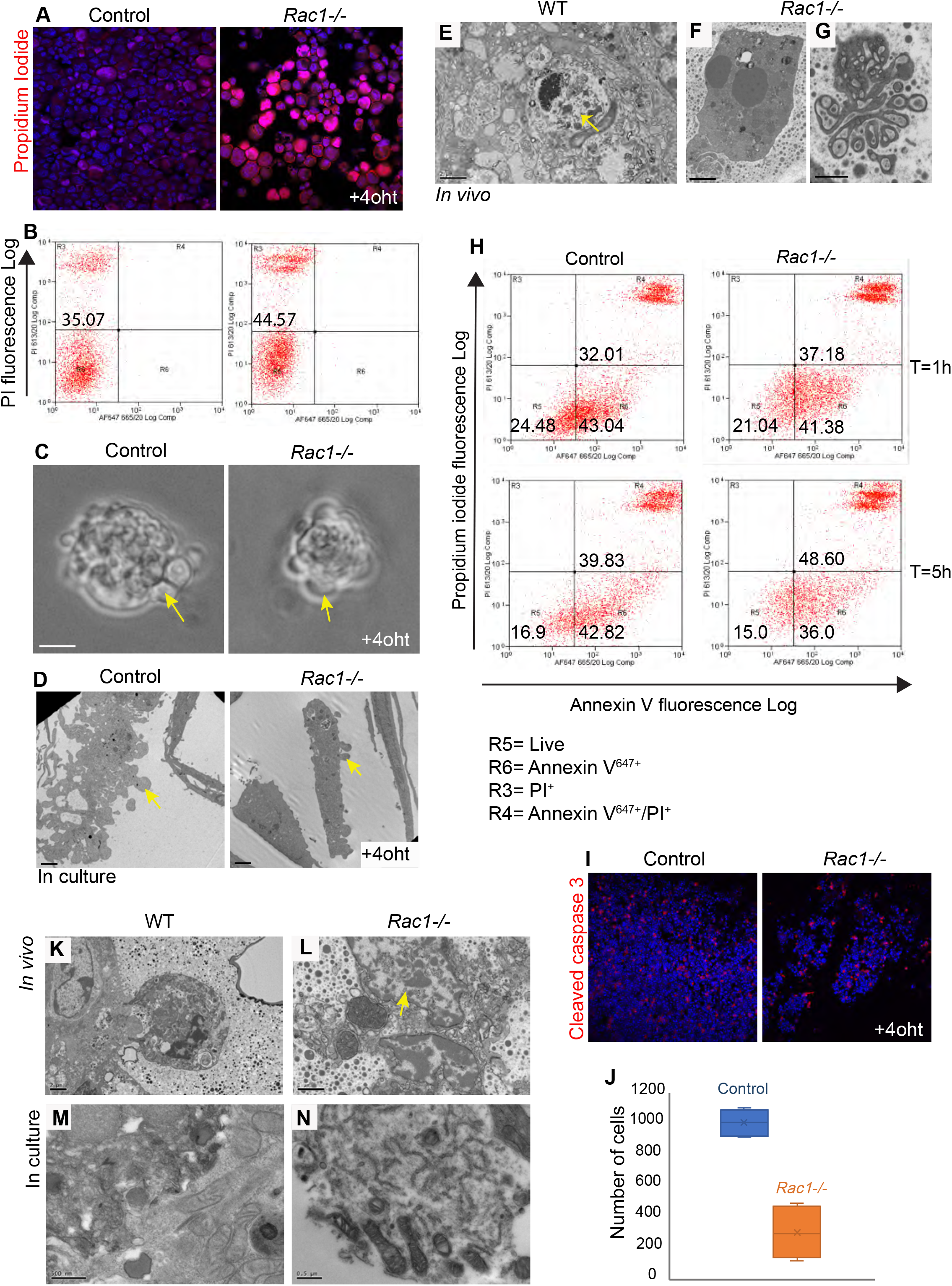
Cell death proceeds faster without Rac1. (A,B) Increased propidium iodide uptake in *Rac1-/-* cells induced to die through anoikis in culture; (A) fluorescent image and (B) quantification with FACS. (C,D) Apoptotic blebs in WT and *Rac1-/-* dead cells in culture detected using light microscopy (C) and EM (D). Arrows: membrane blebs. Bar; (C) 5μm (D) 2μm. (E-G) EM images of (E) WT and (F,G) *Rac1-/-* tissues in vivo show apoptotic cells with nuclear pyknosis (E,F) and membrane blebbing (G). Arrow; apoptotic cells are engulfed by the alveolar epithelium in WT glands (E). Bar; (E,F) 2μm, (G) 1μm. (H) Annexin V-^647+^ and PI^+^ co-labelled WT and *Rac1-/-* cells quantified by flow sorting following 1h and 5h in suspension show more cells proceed to late stage apoptosis/necrosis without Rac1. (I) WT and *Rac1-/-* cells induced to die through anoikis for 8h show reduced numbers of viable cells without Rac1. Cleaved caspase-3 was used to stain apoptotic cells. (J) Quantification of (I) showing total number of WT and *Rac1-/-* cells. Error bars; +/- SEM of n=4. (K-N) EM images of WT and *Rac1-/-* in vivo tissues (K,L) and primary cultures (M,N) show cell necrosis without nuclear pyknosis and organelle release in *Rac1-/-* cells (L,N). In contrast dying WT cells (K,M) have an intact cell membrane. Arrow: Nucleus released from necrotic cell without condensation. Bar: (K) 2μm, (L) 1μm, (M,N), 0.5μm.

As Annexin V can also bind necrotic cells with ruptured membranes, some of the cells in the AnnexinV/PI fraction may be a result of primary necrosis. To investigate primary necrosis we analysed the nuclei of necrotic cell corpses by electron microscopy. In *Rac1-/-* glands, the nuclei of ruptured cells were not condensed, suggesting they had not entered the apoptotic pathway first, but rather died through primary necrosis (Figure 7L). Organelle spillage and cell necrosis was also detected in *Rac1-/-* primary cultures (Figure 7N). In contrast WT dead cells had intact membranes with nuclear condensation (Figure 7K,M). These studies reveal that Rac1 slows down the process of programmed cell death. Without Rac1, cells die either through primary necrosis or through apoptosis but the apoptotic death proceeds more rapidly than in WT epithelia. Taken together, loss of Rac1 increases cell turnover rates in the involuting mammary gland through both increased progenitor proliferation and accelerated cell death.

### Lactation fails to resume upon pup re-suckling in involuting Rac1-/- glands

To determine the functional consequences of the increased cell turnover in *Rac1-/-* glands, we investigated the reversible phase of the involution process. Nursing WT and *Rac1-/-* dams were separated from the pups for 48h to stimulate the first phase of involution and then reunited for a 24h period. Re-suckling in the WT mammary glands recommenced lactation as detected by an approximate 18-fold increase in *casein 2* gene expression and the alveolar epithelium resumed a lactation morphology (Figure 8A, C). In contrast the *Rac1-/-* glands failed to lactate and the involution morphology persisted with dead cell shedding into the lumen (Figure 8B, C). Whilst a small increase in milk protein gene expression was detected in re-suckled *Rac1-/-* glands compared with the involuting *Rac1-/-*, the *casein* gene expression was 18-fold less than the WT fed gland. This compromise in milk protein expression is significantly greater than the 2-fold decrease detected in the first lactation cycle where *Rac1-/-* dams are still able to support pups (9). Together these data indicate that Rac1 is crucial for mammary gland reversibility in phase I of the involution process upon re-suckling. Without Rac1, cell turnover rates increase in early involution but the newly replenished alveoli lose the ability to recommence lactation.

**Figure 8:**
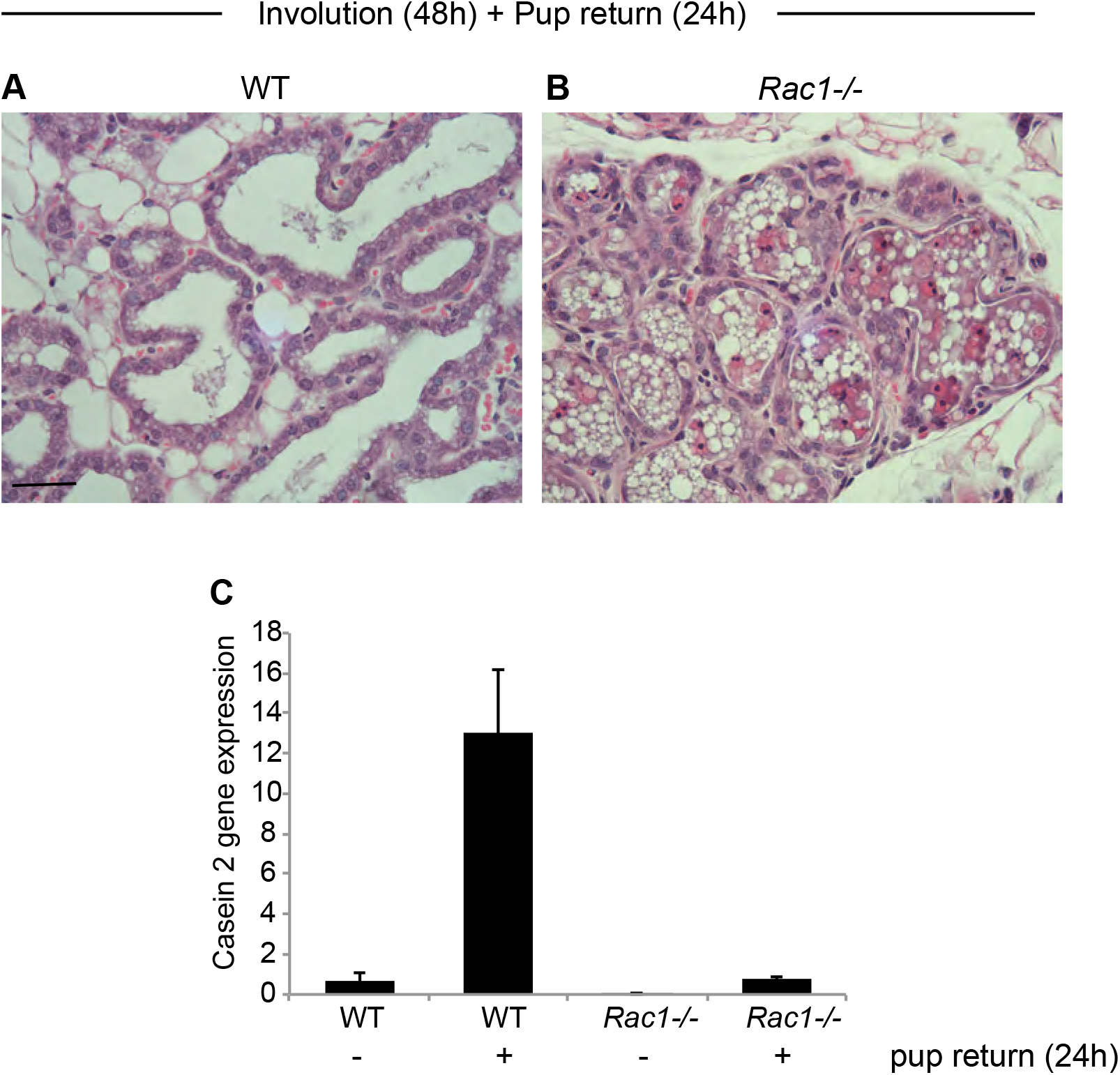
Mammary gland reversibility fails in Rac1-/- involuting glands. (A,B) H+E stain of WT and *Rac1-/-* glands involuted for 48h and then re-suckled for 24h. WT glands (A) resume a lactation morphology while *Rac1-/-* (B) show an involution phenotype. (C) Quantitative RT-PCR shows increased *casein 2* gene expression in re-suckled WT glands but this is reduced by 18-fold in *Rac1-/-* glands. Error bars; +/-SEM of n=3 mice.

## Discussion

Our study reveals that Rac1 acts as a central nexus in controlling the balance of cell death and proliferation within the mammary gland and is critical for mammary gland reversibility in early involution. The mammary gland removes approximately 90% of its tissue weight in post-lactational involution with mass destruction of the milk secreting alveolar units. This extensive regression can only be accomplished if the balance tips towards cell death with reduced proliferation. We have discovered that cell proliferation actively stops in involution and this is a Rac1-mediated process. Removal of Rac1 induces extensive proliferation within the involuting gland. We show the initial delay in alveolar regression is a result of compensatory cell renewal within alveoli and not a delay in cell death. The newly replaced cells in alveoli, however, have a short lifespan as the mammary alveoli regressed by 4 weeks post-weaning. A previous study also reported a delay in *Rac1-/-* alveolar regression but attributed the effects to delayed cell death (23). In contrast our data show that cell death is not blocked in *Rac1-/-* alveoli in early involution as we have detected numerous cell corpses using both light and electron microscopy.

The involution process is accompanied by inflammatory cell influx to remove residual dead corpses by phagocytosis, increased matrix metalloproteinase activity, extracellular matrix remodeling with subsequent release of various morphogens (13, 24). It is well established that pro-inflammatory signals invoke stem cell proliferation in several disease models including psoriasis and various cancers. The involution microenvironment is known to support breast cancer growth in tumour mouse models and promote postpartum breast cancer in women (24, 25). We have made the important discovery that one mechanism by which the postpartum involuting mammary gland protects itself from inflammation-induced proliferation is through the Rac1 GTPase. Consequently healthy mammary epithelial progenitor cells resist proliferating in a tumour promoting microenvironment. Loss of Rac1 increases pro-inflammatory signals in the mammary gland, thereby stimulating cell proliferation. Interestingly in the skin, loss of Rac1 also causes stem cell release through activation of c-myc, although this study did not investigate inflammatory responses (26). Whether Rac1 promotes or suppresses cell proliferation appears to be dependent on the microenvironmental context. Rac1 is linked to stem cell renewal and cell cycle progression in mammary epithelia and in numerous other models (17, 19–22). We have now shown that Rac1 genetic deletion perturbs proliferation in purified epithelial organoid cultures void of impending inflammatory cells. This suggests both cell autonomous and non-autonomous regulation of proliferation by Rac1 depending on the environmental context. In addition to heightened inflammatory signals, stagnant milk in *Rac1-/-* mammary gland structures might cause stretch-induced proliferation. Indeed ductal and alveolar bloating is severe in *Rac1-/-* glands because of defective clearance by MEC phagocytes.

Upstream of Rac1, β1-integrin has been linked to stem cell renewal, cell cycle progression and lactational differentiation in mammary epithelia (16, 18, 19, 27, 28). Here we demonstrate that in involution, Rac1 controls alveolar regression independently of β1-integrin. This suggests distinct upstream wiring allows Rac1 to perform multifaceted roles within the mammary gland. The Rac guanine nucleotide exchange factors (GEFs) ELMO and Dock180 also show delayed alveolar regression in involution but the receptor that activates these GEFs remains to be identified (23).

Our data show that alveolar epithelial cells die through distinct mechanisms with and without Rac1. Multiple cell death mechanisms have been reported in mammary gland involution, including autophagy and lysosomal leakiness linked to milk fat engulfment (6, 7, 29). We discovered that in the absence of Rac1, cell death with autophagy is impaired, milk phagocytosis is impaired (9) with a concomitant lack of detectable lysosomes by EM and down-regulation of several lysosomal genes. Despite these deficiencies, *Rac1-/-* cells still die through both apoptotic and necrotic mechanisms and there is no delay in cell death. These alternate routes ensure a protective redundancy that enables cell death to proceed. Of interest is that in the autophagy defective *Beclin 1-/-* mammary glands, Rac1 activation is perturbed, which suggests a regulatory feedback loop (7).

We have discovered that in addition to increased proliferation, apoptosis accelerates without Rac1, thereby Rac1 critically functions to limit cell turnover rates in involution. We have identified this as an important self-regulatory mechanism that enables mammary gland reversibility within the first 48h of involution. Despite cell renewal within *Rac1-/-* alveoli in early involution, the replaced cells cannot lactate upon re-suckling. This suggests a mechanism by which existing cells resist cell death to allow reversibility. One such mechanism is autophagy, which might act as a survival mechanism in early involution instead of cell death. Ultimately the replaced cells in *Rac1-/-* alveoli also succumb to death as the gland regressed 4 weeks post-weaning involution. The lactation defect, however, is long-term, as we have previously demonstrated severely defective future lactations in *Rac1-/-* mammary glands (9). Future studies will focus on how perturbing Rac1 alters the luminal stem/progenitor niche leading to defective alveolar lineages and long-term tissue malfunction in successive gestations.

## Methods

### Mice

The *Rac1^fl/fl^ :YFP;WAPiCre^Tg/•^* and *Rac1^fl/fl^;CreER^TM^* mice were as previously described (15). For the *in vivo* analysis, the WAPiCre promoter, which is activated mid-late pregnancy was used for *Rac1^fl/fl^* gene deletion specifically in luminal mammary epithelial cells. *Rosa:LSL:YFP* reporter gene was used to detect Cre-induced recombination of flox alleles. *Rac1^fl/fl^ :YFP* littermates that lacked the *Cre* gene were used as WT controls. The genotypes of offspring were determined by PCR amplification of ear DNA as in (15). *Rac1^fl/fl^:YFP;CreER^TM^* mice were used for inducible deletion of the Rac1 gene in primary cultures. Female mice were mated between 8 and 12 weeks of age. *β1-integrin^fl/fl^:YFP:WAPiCre* mice were generated by crossing *β1-integrin^flfl^* mice; JAX #004605 (30) with *WAPiCre:YFP* mice previously described (9). For involution studies, dams were allowed to nurse litters (normalized to 6-8 pups) for 7-10 days and then pups were weaned to initiate involution. In the involution rescue experiments, breeding trios were set up with a male, an experimental female and a WT surrogate female. Impregnated females were separated from the male, allowed to litter and nurse offspring jointly as above. Experimental dams were separated for 48h to involute, while the surrogates continued feeding the pups. Litters were subsequently reunited with the experimental female for a period of 24h prior to gland harvesting. 3-6 mice per group were analysed for each developmental stage. Mice were housed and maintained according to the UK Home Office guidelines for animal research.

### Histology

Mammary tissue was formalin fixed (4% v/v), paraffin embedded before sectioning (5 μm) and subjected to standard haematoxylin and eosin (H&E) staining. Histology was imaged using 3D Histotec Pannoramic 250 slide scanner and Aperio ImageScope version 12.1 software. Adipocytes were quantified using Fiji/ImageJ adiponectin software plug-in.

Whole mount analysis was performed by spreading inguinal mammary glands on polysine slides and stained with carmine alum as previously described (27). Glands were imaged using a Nikon SMZ18 stereoscope.

### TEM

Involuting mammary tissues were fixed with 4% formaldehyde + 2.5% glutaraldehyde in 0.1M Hepes buffer (pH 7.2) for 1hour. Post-fixed with 1% osmium tetroxide + 1.5% potassium ferrocynaide in 0.1M cacodylate buffer (pH7.2) for 1 hour, in 1% thyocarbohydrazide in water for 20 min, in 2% osmium tetroxide in water for 30 min, followed by 1% uranyl acetate in water for overnight. The next day tissues were stained with Walton lead aspartate for 1 hour at 60°C degree, dehydrated in ethanol series infiltrated with TAAB 812 hard grade resin and polymerized for 24h at 60°C degree. TEM sections were cut with Reichert Ultracut ultramicrotome and observed with FEI Tecnai 12 Biotwin microscope at 100kV accelerating voltage. Images were taken with Gatan Orius SC1000 CCD camera.

### RNA isolation and cDNA synthesis

RNA was isolated from the 4^th^ inguinal mammary gland using Trifast reagent (Peqlab) according to manufacturer’s instructions. 2μg of RNA was used to prepare cDNA using the high capacity RNA to cDNA kit (Invitrogen) according to manufacturer’s instructions.

### Affymetrix gene array

Gene arrays were conducted previously(9), Data accession: E-MTAB-5019 (Array Express) or GSE85188 (GEO). Gene lists were analysed using DAVID, Panther and GSEA web accessible programs.

### Quantitative RT-PCR

QRT-PCR was performed on the Applied Biosystems 7900HT using the following Taqman probes (Applied Biosciences; Life Technologies): Krt18 (Mm01601704_g1), MAPKI (Mm00442479_m1) as house-keeping genes, CCL2 (Mm00441242_m1), CCL7 (Mm00443113_m1), Csn2 (Mm04207885_m1). Thermal cycling conditions were: UNG start at 50°C (2min), 95°C (10min) and then 40 cycles of 95°C (15 sec) followed by annealing at 60°C (1 min).

### Primary cell culture and gene deletion

Primary MECs were harvested from either 15.5-17.5 day pregnant mice for alveolar cultures or 12 week old virgin mice for ductal cultures and cultured as described in (31).

Cells were plated onto Collagen 1 for monolayer cultures or basement membrane-matrix (Matrigel; BD Biosciences) to form acini and cultured in growth media (Ham’s F12 medium (Sigma) containing 5 μg/ml insulin, 1 μg/ml hydrocortisone (Sigma), 3 ng/ml epidermal growth factor (EGF), 10% fetal calf serum (Biowittaker), 50 U/ml Penicillin/Streptomycin, 0.25 μg/ml fungizone and 50 μg/ml gentamycin). For ductal branching cells were cultured in branching medium (DMEM-F12 medium (Lonza) containing insulin-transferrin-selenium G supplement (100x), 2nM hb-fibroblast growth factor (Sigma cat: F0291) and 50 U/ml Penicillin/Streptomycin). For inducible Rac1 gene ablation, MEC cultures were prepared from *Rac1^fl/fl^;CreER^TM^* mice, and treated with 100 nM 4-hydroxytamoxifen dissolved in ethanol. For proliferation assays, 10μM EdU was added for 2h to either day 2 or day 3 cultures and developed using the Click-IT EdU Alexa Fluor imaging kit (Life Technologies).

### Anoikis assays and FACS

Apoptotic MECs (-/+4OHT) were prepared in suspension culture for either 1h or 5h, in serum-free media (DMEM-F12 containing 2.5μg/ml insulin). Cells were labelled with propidium iodide and/or Alexa Flour 647-Annexin V (Biolegend) according to manufacturer’s instructions and analysed by flow cytometry or cytospun onto polysine slides, fixed and counterstained with hoechst for micrographs. Some cytospun suspension cultures were stained with cleaved caspase-3 antibody.

### Immunostaining

Expression and distribution of various proteins were visualised by indirect immunofluorescence. Cells were fixed for 10 min in PBS / 4% (w/v) paraformaldehyde, and permeabilised for 7 min using PBS / 0.2% (v/v) Triton X100. Non-specific sites were blocked with PBS / 10% goat serum (1h, RT) prior to incubation with antibodies diluted in PBS / 5% goat serum (1h, RT, each). EdU was detected using the Click-iT EdU Alexa Fluor 647 imaging kit (Life Technologies) and nuclei were stained using 4 μg/ml Hoechst 33258 (Sigma) for 5 min at RT. Cells were washed in PBS before mounting in prolong gold antifade (Molecular Probes). Images were collected on a Nikon A1 confocal or a Leica TCS SP5 AOBS inverted confocal as previously described (15). Non-biased cell counts were performed by concealing the identity of each slide.

Immunostaining of mammary tissue was performed on paraffin-embedded tissue (5μm) or cryosections (10μm) and luminal surface was detected with wheat germ agglutinin-488, or −647 (Invitrogen, #W11261, #W32466) and imaged using confocal microscopy. Primary antibodies used for immunofluorescence are indicated in Supplementary table 2. Secondary antibodies were conjugated to Cy2, Alexa-488, Rhodamine-RX, Cy5, Alexa-647 (Jackson Immunoresearch) and horse-radish peroxidase.

### Protein analysis

Proteins were extracted as in (16). Equal amounts of proteins were used and equivalent loading assessed by referral to controls, such as Calnexin (Bioquote SPA-860) or E-cadherin (Cell Signalling #610182). Primary antibodies used for immunoblotting are indicated in Supplementary table 2. ImageJ was used to quantify bands.

## Abbreviations

WT: WT
MEC: mammary epithelial cell

## Author contribution

NA conceived ideas, performed most of the experiments and wrote the manuscript. MF, RE and MK contributed experiments, AM performed the electron microscopy.

## Acknowledgements

We thank Karen-Piper Hanley (University of Manchester) for the *β1-integrin^flfl^* mice. This work was supported by the Thomas, Berry and Simpson fellowship (N.A), Wellcome Trust #200588/Z/16/Z (N.A) and Medical Research Council; MR/P028411/1 (N.A).

The authors declare no financial or conflicts of interest related to this manuscript.

